# Specialized Pro-Resolving Mediator loaded Extracellular Vesicles Mitigate Pulmonary Inflammation

**DOI:** 10.1101/2025.04.09.648009

**Authors:** Manjula Karpurapu, Jiasheng Yan, Sangwoon Chung, Sonal R Pannu, Narasimham Parinandi, Evgeny Berdyshev, Liwen Zhang, John W Christman

## Abstract

Specialized pro-resolving mediators (SPMs), including lipoxins derived from arachidonic acid and resolvins, protectins, and maresins derived from docosahexaenoic acid (DHA) and eicosapentaenoic acid (EPA), orchestrate the active resolution of inflammation. These SPMs are biosynthesized through the coordinated interaction of various cells in a process known as transcellular biosynthesis, involving the sequential action of cyclooxygenase-2 (COX-2), 5-lipoxygenase (5-LOX), 12-lipoxygenase (12-LOX), and/or 15-lipoxygenase (15-LOX) enzymes. Additionally, Aspirin-triggered Resolvins are produced by acetylated COX-2, along with various lipoxygenases. Although SPMs regulate various cellular processes to actively resolve inflammation, their in vivo levels are typically low. To address this limitation, we engineered a multigene expression vector that co-expresses COX-2, 5-LOX, and 15-LOX, potentiating the synthesis of various SPMs. HEK293T cells transfected with this vector and cultured with fatty acid-free BSA-complexed DHA, EPA, and aspirin, successfully mimicked both transcellular and aspirin-triggered biosynthesis of Resolvins. These Resolvins are packaged into extracellular vesicles, which significantly inhibited neutrophil adhesion to endothelial cells, preserved endothelial monolayer barrier integrity, suppressed NF-κB reporter activity, and enhanced macrophage efferocytosis *in vitro*. Notably, post-injury administration of Resolvin-loaded EVs mitigated pulmonary inflammation in LPS-treated mice without causing systemic or pulmonary toxicity. In summary, we report a novel cell-based platform for generating Resolvin-loaded EVs that mitigate pulmonary inflammation in mouse models, underscoring their potential for treating other acute inflammatory diseases.

## Introduction

Acute inflammation is a critical response of the body to injury or infection, characterized by the rapid activation of the innate immune system and the release of pro-inflammatory mediators. This process, while essential for tissue repair and pathogen defense, can become dysregulated and lead to tissue damage if unresolved. Cellular injury and inflammation triggers release of arachidonic acid, an ω-6 polyunsaturated fatty acid (PUFA), from the phospholipids of the cell membrane catalyzed by different phospholipase A2 (PLA2) members^1,2^. Arachidonic acid released during inflammation serves as a precursor for different bioactive eicosanoids. Arachidonic acid is further metabolized by the COX pathway enzymes COX-1 and COX-2, generating prostanoids (PGE2, PGD2, PGF2α, PGI2), and thromboxanes (TXA2); LOX pathway enzymes (5-LOX, 12-LOX, and 15-LOX, 12/15-LOX in mice), producing leukotrienes (LTA4, LTB4, LTC4, LTD4, LTE4), lipoxins (LXA4, LXB4), and -Hydroperoxyeicosatetraenoic acids (12- or 15-HPETE, further reduced to hydroxyeicosatetraenoic acids-HETES); and CYP450 pathway by CYP450 epoxygenase and CYP450 ω-hydroxylase, generating epoxyeicosatrienoic acid (EETs) and (ω-HETEs)^3–5^. Specific members of prostanoids and HETEs are linked to inflammation as they mediate PMN chemotaxis, vascular permeability and oxidative stress^6–8^. In contrast, pro-resolving lipoxins biosynthesized from arachidonic acid; resolvins, protectins and maresins biosynthesized from the ω-3 PUFAs (EPA and DHA), collectively referred to as SPMs, are implicated in the resolution of inflammation and restoration of tissue homeostasis^9–12^. Interestingly, certain lipid mediators including LTB4, Lipoxins, resolvins and maresins, are produced through a coordinated interaction between different cells, a process known as ‘transcellular synthesis’ ^13–16^. Both the D and E series resolvins are typically synthesized through the sequential action of COX2, 5-LOX,12-LOX and 15-LOX enzymes, which require coordinated interactions between different cell types, including endothelial, epithelial cells, monocytes/macrophages and granulocytes. In addition, aspirin-acetylated COX-2 converts EPA to 18R-hydroperoxyeicosapentaenoic acid (18R-HEPE) in vascular endothelial cells, which is then transformed by activated polymorphonuclear leukocytes (PMNs) into bioactive Aspirin triggered Resolvin E (AT-Resolvin E) members^17^. Similarly, DHA is converted to 17R-hydroxydocosahexaenoic acid (17R-HDHA) and AT-Resolvin D series members. SPMs were shown to promote active resolution of inflammation by inhibiting important cellular events of inflammation including trans-endothelial and trans-epithelial migration of neutrophils, increase in macrophage efferocytosis to clear apoptotic cells, anti-inflammatory cytokine IL-10 release; inhibit proinflammatory cytokine production and inflammasome activation^18–22^.

Extracellular vesicles (EVs) are lipid bilayer membrane-bound structures released from most living cells into the extracellular space which originate from two main sources: small EVs are derived from the cellular endosomal system, and microvesicles (MVs) are released directly from the plasma membrane^23–25^. Notably, EVs transfer proteins, micro-RNAs, lipids, and various membrane proteins, including signal transduction receptors to the recipient cells^26–30^. Different proteins and micro-RNAs (miRs) transported by EVs were shown to regulate inflammatory signaling pathways in pulmonary diseases, including ARDS^31–34^. Our group, for the first time, demonstrated that the small EVs (50-150 nm) from mouse bronchoalveolar lavage fluid (BALF) carried about 140 different lipid mediators, released by the COX and LOX enzymes from both ω-6 and ω-3 PUFAs^35, 36^. Furthermore, the amount of lipids packaged in EVs is significantly higher than that of EV-depleted BALF. Although found in significantly low amounts, Lipoxin A4, Resolvin D1 and D6 were also detected in BALF EVs, indicating the propensity of pulmonary cells to package SPMs into EVs^35^. Current research suggests that these lipid mediators have short half-life, act close to the site of synthesis in a transient autocrine or paracrine manner or in circulation/pulmonary edema fluid after their generation. On the other hand, recent studies from our group and others demonstrate that lipids transported by small EVs are stable due to their packaging within a saturated lipid-rich bilayer membrane^35–40^. Therefore, it is likely that EVs increase the stability of the biological cargo, which mediates intercellular communication. Additionally, during inflammatory conditions, the compromised intercellular barrier enhances the uptake of EVs by distant cells^41^.

In the current manuscript, we report an innovative strategy to package and deliver different Resolvin D and E members directly to lungs, to mitigate acute inflammation-induced by LPS. In brief, we co-expressed COX2 and LOX proteins to harness transcellular and aspirin triggered-resolvin synthesis and leveraged the intrinsic ability of EVs to encapsulate various SPMs, in two distinct approaches. In the first approach, we co-expressed human COX2 and 5-LOX proteins together in HEK293T cells. The transfected HEK293T cells grown in serum-free medium, supplemented with 5 µM each of DHA and EPA (complexed to fatty acid free BSA) and 1 mM aspirin released EVs that contained Resolvin D1 and E1 (referred to as Resolvin D1E1 EVs). However, resolvin loading efficiency in this method requires equal transfection efficiency of both COX2 and 5-LOX plasmids. To reduce the inconsistencies associated with the use of two different plasmids, in the second method, we custom synthesized a multigene expression vector *pRP_PTGS2_ALOX5_ALOX15* (VectorBuilder Inc. Chicago IL) that synthesizes PTGS2 (COX2), ALOX-5 (5-LOX) and ALOX-15 (15-LOX) simultaneously, which increases the number of various Resolvins released by these proteins. The COX2, 5-LOX, and 15-LOX proteins were separated by P2A and T2A viral sequences for independent protein production. Notably, EVs released from HEK293T cells transfected with the multigene expression vector, grown in presence of 10 µM each of DHA, EPA and aspirin mix are packaged with a wider range of Resolvin D and E members (referred to as Resolvin EVs). Both the Resolvin D1E1 EVs and Resolvin EVs contained resolvins each at concentrations exceeding physiological levels in mice by several thousand-fold. EVs generated using both methods exhibited significant pro-resolution activity *in vitro* across PMVEC, neutrophils and macrophages. Compared to Control EVs the Resolvin EVs significantly mitigated acute lung injury in LPS-treated mouse models. Our findings introduce a novel strategy to load EVs with several resolvin members that mitigate pulmonary inflammation in mouse models, underscoring their potential for treating other acute inflammatory diseases.

## Results

### Selective loading of Resolvin D1 and E1 into EVs (RvD1E1-EVs)

Biosynthesis of Resolvin D and E members from DHA and EPA requires coordinated action of acetylated COX2 and 5-LOX enzymes (Figure 1a). To selectively steer the cells towards Resolvin synthesis, HEK293T cells were transfected with human pcDNA3.1_COX-2 and pcDNA3.1_5-LOX plasmids and cultured in the presence of DHA and EPA, 5 µM each complexed to fatty acid-free bovine serum albumin (BSA) and 1 mM aspirin (Figure 1b). Size of EVs isolated from the culture medium was in the expected size range of 80-120 nM, assayed by nanoparticle tracking analysis (Figure 1c, 1d). Control (pcDNA3.1) or COX2 and 5-LOX expression plasmid transfected HEK293T cells and EVs were analyzed for expression of COX2 and 5-LOX proteins. Both COX2 and 5LOX were over-expressed in cell lysates, but undetectable in EVs and control plasmid transfected cell lysates (Figure 1e). Calnexin was enriched in cell lysates but not detected in EVs. In addition, CD9 was enriched in EVs, but undetectable in the cell lysates because the amount of protein loaded was minimal to be able to match the low protein levels present in EVs. Similarly, COX2 and 5-LOX mRNA were significantly enriched in the cells transfected with COX2 and 5-LOX plasmids but undetectable in EVs and control vector transfected HEK293T cells (Figure 1f). It is important to note that the absence of COX2 and 5-LOX mRNAs in Resolvin D1E1 EVs rules out their transfer to recipient cells. EVs from pcDNA Control vector transfected cells grown in 0.001% ethanol (solvent control) or DHA, EPA, Aspirin mix showed no detectable ResolvinD1 and E1 (Table 1). Similarly, pcDNA_COX2 and pcDNA_5-LOX transfected cells grown in the presence of ethanol released EVs (pcDNA_COX2+pcDNA_5-LOX EVs) with no detectable Resolvins. Notably, EVs from COX2 and 5-LOX transfected cells grown in presence of DHA, EPA and aspirin (referred to as RvD1E1 EVs) contained about 62 ng Resolvin D1 and 12.9 ng Resolvin E1/mg EV protein, analyzed by LC-MS/MS (Table 1). Furthermore, the RvD1E1EVs showed no detectable eicosanoids including PGD2, LTE4, 12-HETE, 15-HETE, 13HODE, LTE4 and Lipoxin A4 derived from AA. However, significant levels of PGE2 was detected in RvD1E1 EVs. PGE2 acts both as pro-inflammatory and pro-resolution lipid, depending on the local cellular signaling environment. It is important to note that although we detected a wide array of LMs, COX2 and 5-LOX mRNAs were not packaged into EVs.

**Figure 1.**
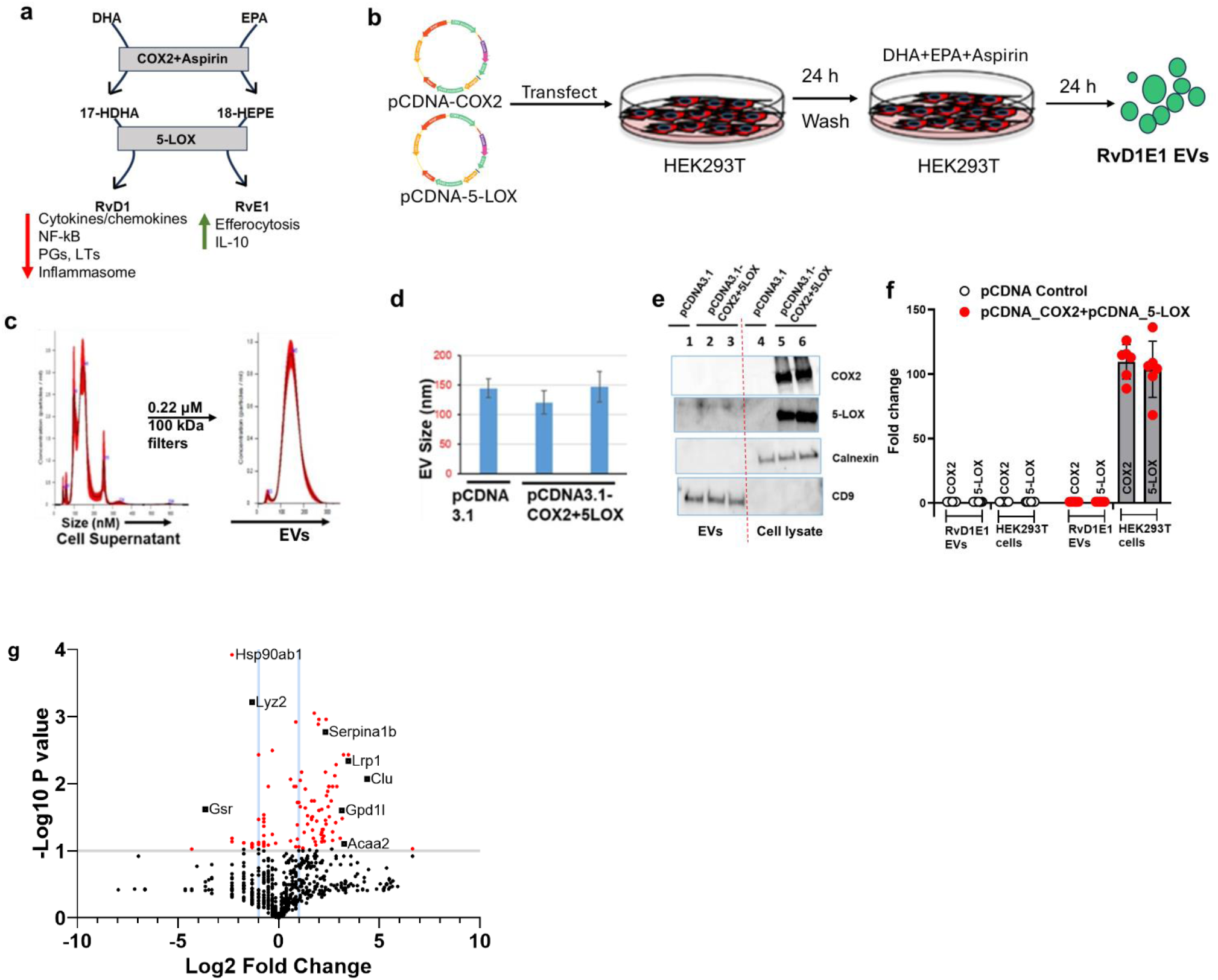
Characterization of Resolvin D1 and E1-loaded EVs from HEK293T cell supernatants. a) Schematic illustrating the aspirin-triggered biosynthetic pathway of Resolvin D1 and E1. b) Cell-based method for overexpressing COX2 and 5-LOX to load Resolvin D1 and E1 into EVs. c-d) EV size is not significantly different between control and RvD1E1 EVs, as determined by nanoparticle tracking analysis (NTA). e) Immunoblotting for EV and cell-specific marker proteins. CD9 was detected in EVs, while Calnexin was absent, and only HEK293T cells transfected with COX2 and 5-LOX vectors showed Calnexin, COX2, and 5-LOX proteins. f) COX2 and 5-LOX mRNA were overexpressed in the pcDNA3.1_COX2 and pcDNA_5LOX transfected HEK293T cells but were undetectable in all EV groups and the Control pcDNA3.1 vector transfected cells, analyzed by qRT-PCR. g) BALF EV proteome analyzed by Orbitrap mass spectrometer showing the lipid metabolism-related proteins.

**Table 1.** Detection of pro-resolution lipid mediators packaged in EVs by LC-MS/MS. EVs released from HEK293T cells transfected with different plasmid vectors grown in 0.001% ethanol (vehicle control) show no detectable resolvin D1 and E1. RvD1E1EVs from pcDNA_COX2 and pcDNA_5-LOX transfected HEK293T cells grown in the presence of DHA, EPA and aspirin show significantly elevated levels of Resolvin D1, E1 and PGE2.

| Lipid mediator | pcDNA Control<br>Vector + Ethanol | pcDNA Control<br>Vector +<br>DHA+EPA+Aspirin | pcDNA3_COX2+<br>pcDNA_5 LOX+<br>Ethanol | pcDNA_COX2+<br>pcDNA_5-LOX+<br>DHA+EPA+Aspirin<br>(RvD1E1 EVs) |
| --- | --- | --- | --- | --- |
| PGE2 | - | - | - | 6.9 ng/mg EV protein |
| PGD2 | - | - | - | - |
| 12-HETE | - | - | - | - |
| 15-HETE | - | - | - | - |
| 13-HODE | - | - | - | - |
| LTE4 | - | - | - | - |
| LXA4 | - | - | - | - |
| RvD1 | - | - | - | 61.87 ng/mg EV protein |
| RvE1 | - | - | - | 12.97 ng/mg EV protein |

### Absence of PUFA catalytic proteins in EVs

Although EVs from transfected HEK293T cells showed no detectable COX2 or 5-LOX proteins or mRNA, we sought to confirm whether the same holds true under physiological conditions. We analyzed the EVs from mouse bronchoalveolar fluid (BALF) using high-throughput LC-MS/MS to determine whether they transport the COX2, 5-LOX and 15-LOX catalytic proteins under physiological conditions. BALF EVs from control and LPS-treated mice contained about 1200 different proteins, identified using Scaffold 4.0v software (Figure 1g, supplementary table 1). Majority of these proteins were present in significantly low amounts and not statistically different between LPS and control groups. However, compared to BALF EVs from control mice, EVs from LPS treated mice showed an increased abundance of around 40 proteins, which regulate diverse cellular functions. These include Serpin family members of protease inhibitors, heat shock proteins, integrins, histone proteins and antimicrobial or bactericidal proteins like Lysozyme 2 and clusterin (Figure 1g). When specifically analyzed for lipid-metabolism related proteins, BALF EVs after LPS treatment contained elevated levels of-lipid transport proteins such as Clusterin or Apolipoprotein J (Clu) and Phospholipid transport protein (Pltp); glycerol-3-phosphate dehydrogenase 1 (Gpd1) which regulates the cellular NAD+/NADH potential and glycerophospholipid metabolism; low-density lipoprotein receptor protein 1 (Lrp1) that plays role in lipid homeostasis, and clearance of apoptotic cells; Acetyl-CoA Acyltransferase (ACAA2-regulates Linoleic acid and alpha-linolenic acid metabolisms), apolipoproteins (binds and transport lipids), and fatty acid synthase (FSN) proteins. It is important to note that although we detected a wide array of LMs, the catalytic proteins that generated the LMs (COX2, 5-LOX, 15-LOX) were absent, indicating their intrinsic exclusion from BALF EVs.

### RvD1E1 EVs mitigate *in vitro* cellular inflammation

We determined whether the RvD1E1 EVs alter the recipient cell function *in vitro* assays using a HL60 derived neutrophils, a somatic hybrid endothelial (EAhy926) and mouse macrophage NF-kB luciferase reporter cell lines.

## Pro-resolving activity of RvD1E1 EVs

### Adhesion of HL60-derived neutrophils to activated endothelial cells

The adhesion of PMNs to activated endothelial cells is a critical event in pulmonary inflammation, driven by adhesion molecules that promote PMN recruitment, inflammation, and tissue damage. We assayed the cellular activity of RvD1E1EVs on PMN adhesion to activated endothelial cells, EAhy926 cells grown on 96 well-black polystyrene culture plates were activated with 50 ng/mL of TNFα for 8h. Following this, PKH26 labeled neutrophils were added to the EAhy926 cells and after 1h, Control EVs or RevD1E1EVs were added separately and incubated for a further 15 h. After 16 h cells were washed, and the relative fluorescence (RFU) was measured at 530/570 nm (timeline shown in Figure 2a). We observed increased adherence of PKH26 labelled neutrophils to activated EAhy926 cells in Control EV treatments (Figure 2b). In contrast, RevD1E EVs significantly decreased the neutrophil adherence to EAhy926 cells, indicating their pro-resolution effect. The biological function of RvD1E1 EVs was further confirmed by pre-treating cells with Nystatin and cytochalasin D, which inhibit EV uptake by inhibiting caveolin based and actin polymerization-based internalization. Both Nystatin and Cytochalasin D restored the adherence of HL60 differentiated neutrophils to EAhy926 cells (Figure 2c). EVs from control vector transfected HEK293T cells had not altered the neutrophil adhesion to EAhy926 cells pretreated with Nystatin and Cytochalasin D.

**Figure 2.**
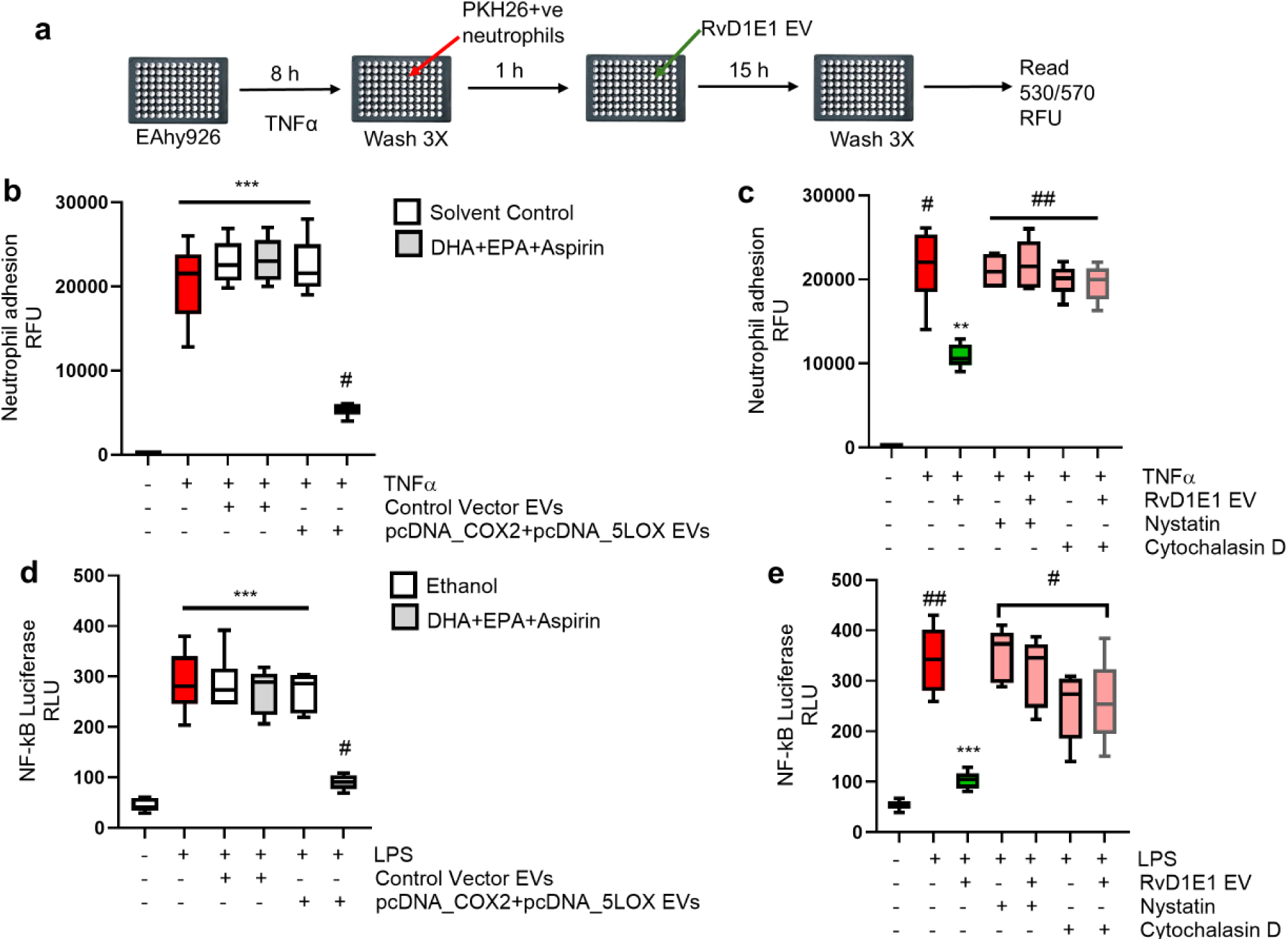
RvD1E1 EVs attenuate neutrophil adhesion and NF-kB activation in *in vitro* cell assays. a) Timeline of the neutrophil adhesion assay. b) RvD1E1 EVs, but not control EVs, reduce the adhesion of HL-60-derived neutrophils to EAhy926 endothelial cells. c) Pretreatment of EAhy926 cells with EV uptake inhibitors, Nystatin and Cytochalasin D, restores neutrophil adhesion to EAhy926 cells in co-culture with RvD1E1 EVs. d) LPS-induced NF-κB luciferase activity is attenuated by RvD1E1 EVs in NF-κB reporter macrophages. e) Pretreatment of NF-κB reporter macrophages with EV uptake inhibitors, Nystatin and Cytochalasin D, restores LPS-induced NF-κB luciferase activity. # p ≤ 0.05, ## p ≤ 0.01, *** p ≤ 0.001.

### LPS-induced NF-kB activation in macrophages

NF-kB plays a vital role in regulating the expression of pro-inflammatory cytokines, chemokines, and adhesion molecules, which contribute to immune cell recruitment, vascular permeability, and tissue damage, all of which are key features of pulmonary inflammation. We measured NF-κB activation in LPS-stimulated NF-kB luciferase reporter macrophages to determine the NF-κB inflammatory signaling in response to RvD1E1 EV treatment. In brief, NF-KB reporter macrophages were stimulated with LPS (50 ng/ml for 30 min) and treated with control EVs or RvD1E1EVs separately, and incubated for an additional 3 h. After 3h, the treated cells were washed, lysed in cell lysis buffer and used to measure luciferase activity using luciferin substrate. As expected, LPS increased NF-kB luciferase reporter activity in Control EV treatments which is a measure of inflammation (Figure 2d). In contrast, RvD1E1 EV treatment significantly decreased NF-kB reporter activity. The cellular activity of EVs was further confirmed by including EV uptake inhibitors Nystatin and cytochalasin D in these assays. The inhibition of RvD1E1EV uptake by Nystatin and Cytochalasin D restored the NF-KB luciferase reporter activity in macrophages confirming the pro-resolution activity of RvD1E1 EVs (Figure 2e).

### Improving the SPM profile of EVs

Since the RvD1E1EV generation method requires successful co-expression of COX2 and 5-LOX using two different plasmids, we sought to improve this approach further. To package additional members of resolvins into EVs, co-expression of 15-LOX along with COX2 and 5-LOX broadens the range of resolvins, as these three enzymes collectively generate a wider range of resolvins. After several iterations, we synthesized a novel multigene expression plasmid *pRP_PTGS2_ALOX5_ALOX15,* capable of co-expressing codon optimized human 15-LOX along with COX2 and 5-LOX proteins (Figure 3a). EVs isolated from HEK293T cells transfected with Control pRP plasmid or pRP_PTGS2_ALOX5_ALOX15 and grown in the presence of DHA, EPA and aspirin EVs were isolated and analyzed. Control pRP plasmid transfected cell lysates showed no detectable COX2, 5-LOX and 15-LOX proteins or mRNA (Figure 3b, 3c). In contrast, pRP_PTGS2_ALOX5_ALOX15 transfected cells showed significantly higher levels of COX2, 5-LOX and 15-LOX proteins and mRNA. LC-MS analysis of EVs from multigene expression plasmid transfected HEK293T cells grown in presence of DHA, EPA and aspirin, referred as Resolvin-EVs, showed significantly higher levels of Resolvin D and AT-Resolvin D members (RvD1, RvD2, RvD5, RvD6, AT-RvD1, AT-RvD5, AT-RvD6) and (RvE1, RvE2) (Figure 3d, 3e). More importantly, we detected elevated resolvin D and E precursors (17-HDHA, 18-HEPE) packaged in Resolvin EVs, in addition to 5-HEPE and 15-HEPE, that possess pro-resolution activity (Figure 3f). Total cell lysates from HEK293T transfected cells with multigene expression vector gown in DHA, EPA, Aspirin also showed all the different Resolvin members listed above in comparable concentrations (data not shown). However, the cell lysates were not analyzed further, as they contain overexpressed COX2, 5-LOX and 15-LOX proteins in addition to several other cytoplasmic and nuclear proteins.

**Figure 3.**
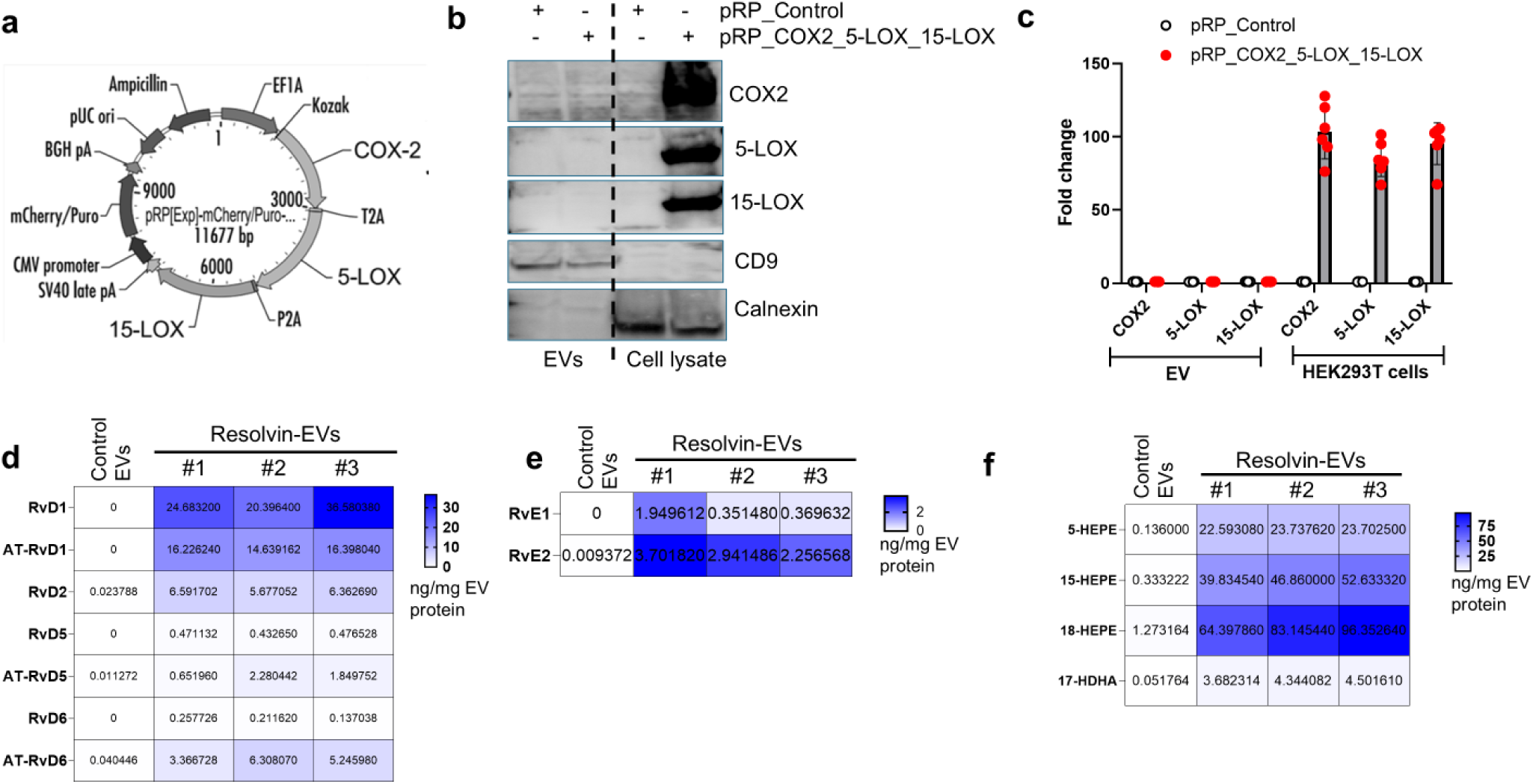
Cell-based system to generate EVs that are loaded with different Resolvin D, Resolvin E members and their precursors. a) Vector map of the custom-synthesized multigene overexpression plasmid designed to co-express COX2, 5-LOX, and 15-LOX proteins, separated by P2A and T2A sequences. b) HEK293T cells transfected with the pRP_PTGS2_ALOX5_ALOX15 plasmid co-express COX2, 5-LOX, and 15-LOX proteins, which are absent in Resolvin-EVs and pRP_Control vector-transfected cells. c) HEK293T cells and EVs transfected with the pRP_Control plasmid or EVs from pRP_PTGS2_ALOX5_ALOX15 plasmid-transfected cells show no detectable transcripts of COX2, 5-LOX, and 15-LOX. Only HEK293T transfected cells contain significant amounts of COX2, 5-LOX and 15-LOX mRNA. Resolvin-EVs contain significant amounts of d) Resolvin D and AT-Resolvin D members; e) Resolvin E1 and E2; and f) resolvin precursors, 18-HEPE and 17-HDHA, analyzed by LC-MS with Turbo Spray detection method. Each resolvin measurement represents pooled EV samples from three independent experiments (n=9 in total). Control EVs (n=3) show negligible amounts of Resolvins. Heat map of Resolvins plotted using GraphPad Prism v10.0. Scale bar = nanograms of lipid/mg EV protein.

## Pro-resolving activity of Resolvin EVs

In an identical experimental set up described in figure 2, we assayed the pro-resolution activity of Resolvin EVs to mitigate PMN adhesion to EAhy926 endothelial cells. As expected, we observed increased adherence of PKH26 labelled neutrophils to activated EAhy926 cells in Control EV treated cells (Figure 4a). In contrast, Resolvin EVs decreased adhesion of HL60 differentiated neutrophils to endothelial cells significantly. Both Nystatin and Cytochalasin D restored the adherence of HL60 differentiated neutrophils to EAhy926 cells indicating the causal relationship between EV uptake and cell adhesion (Figure 4b). Damaged microvascular endothelial barrier integrity is a critical cellular feature that contributes to protein rich fluid extravasation into alveoli, immune cell infiltration, and tissue damage. We asked if the Resolvin EVs impact the *in vitro* microvascular barrier integrity. Pulmonary microvascular endothelial cell (PMVEC) monolayers were grown to 100% confluence on collagen coated trans well inserts and treated with Thrombin (0.2 u/mL). FITC dextran was added to the transwell inserts and its flux across transwells was measured after 24 h. Thrombin increased flux of FITC dextran from transwell inserts to bottom chamber indicating disrupted PMVEC monolayer integrity (Figure 4c). Notably, Resolvin EVs, added 30 min after Thrombin addition, decreased the FITC dextran flux through transwells. Inhibition of Resolvin EV uptake by Nystatin and Cytochalasin, resulted in significant reduction in PMVEC monolayer barrier integrity (Figure 4d). Similarly, in a parallel experiment, Thrombin decreased expression of VE-Cadherin, critical for integrity of PMVEC adherence junctions and barrier integrity (Figure 4e). Resolvin EVs significantly restored VE-Cadherin expression comparable to control EVs. In a parallel experiment, Resolvin EVs attenuated NF-kB activation in macrophages and in contrast, pretreating cells with EV uptake inhibitors restored the NF-KB reporter activity (Figure 4f, 4g). Similarly, post-treatment of LPS stimulated RAW264.7 macrophages with Resolvin EVs significantly decreased the release of pro-inflammatory cytokines IL6 and TNFα (Figure 4h, 4i). Furthermore, we determined whether Resolvin EVs regulate Efferocytosis activity of macrophages using THP-1 derived macrophages and HL-60 derived apoptotic neutrophils. HL-60 derived neutrophils were labelled PKH26 and treated with Staurosporine to induce apoptosis for 6 hours. THP-1 differentiated macrophages were labeled with CytoTell^TM^ Blue and co-incubated with apoptotic neutrophils (4:1 ratio) for 1 hour followed by addition of control or Resolvin EVs. After 16 hours of total co-incubation of macrophages and apoptotic neutrolphils celle were washed and analyzed on flow cytometer. Notably, Resolvin EVs increased efferocytotic activity of macrophages compared to control EVs (Figure 4j gating strategy, 4k). Based on the strong pro-resolving activity exhibited by Resolvin EVs *in vitro* assays, we extended our studies to determine the potential of Resolvin EVs in mitigating pulmonary inflammation in a mouse model of ALI induced by LPS.

**Figure 4.**
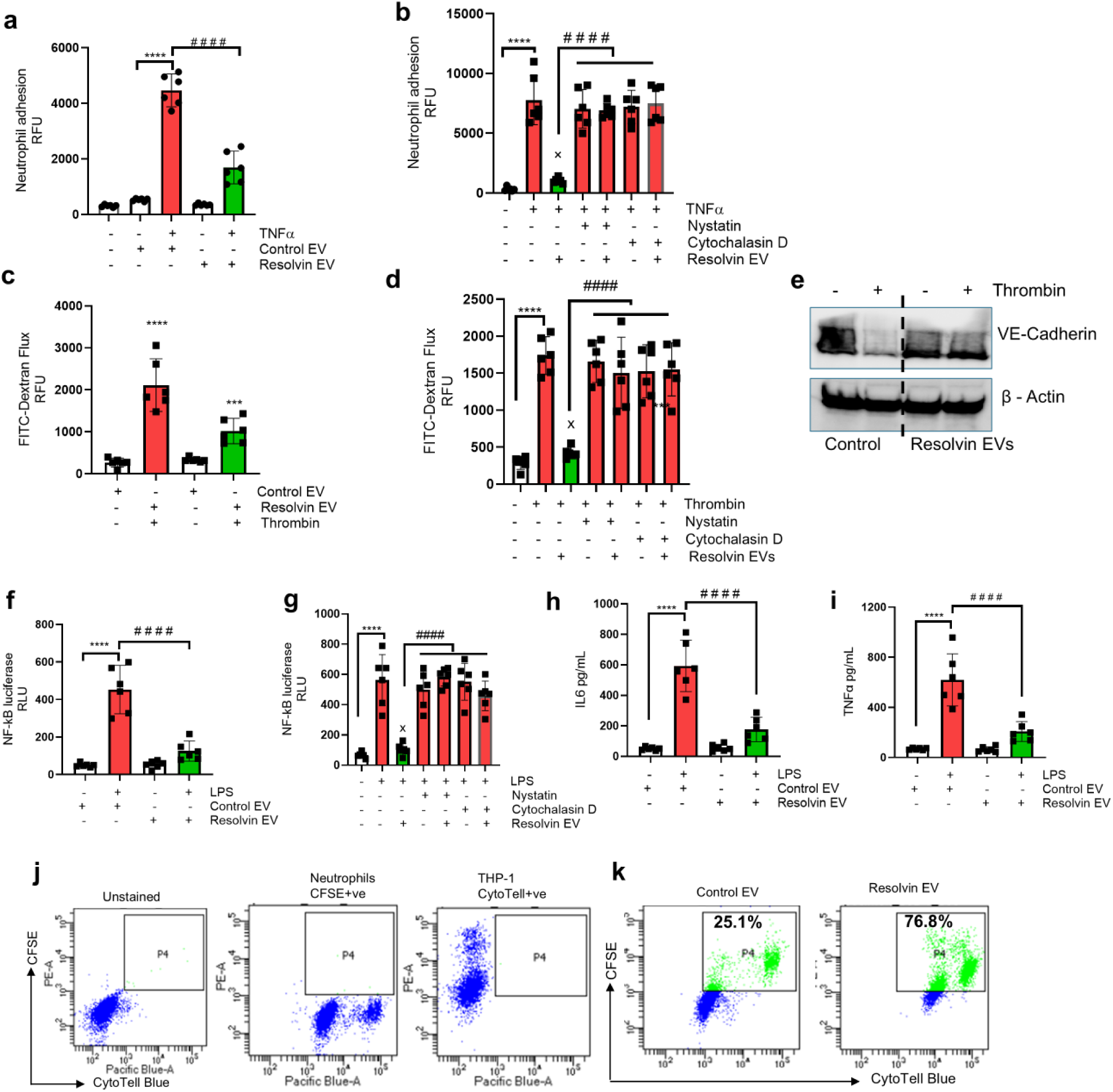
Resolvin-EVs mitigate inflammatory cellular events. a) Compared to control EVs, the Resolvin-EVs decrease the adhesion of HL-60-derived neutrophils to EAhy926 endothelial cells activated by TNFα. b) Pretreatment of EAhy926 cells with EV uptake inhibitors, Nystatin and Cytochalasin D, restores neutrophil adhesion to EAhy926 cells in co-culture with Resolvin-EVs. c) Resolvin-EVs maintain barrier integrity in PMVEC monolayers, as measured by FITC-Dextran flux. d) Pretreatment of PMVECs with EV uptake inhibitors, Nystatin and Cytochalasin D, increases thrombin-induced FITC-Dextran flux through PMVEC monolayers in co-culture with Resolvin-EVs. e) Compared to control EVs, Resolvin-EVs restore the expression of VE-Cadherin, which is downregulated by thrombin treatment in PMVECs. f, g) Resolvin-EVs attenuate the LPS-induced NF-κB reporter activity in macrophages, and inhibition of Resolvin EV uptake by Nystatin and Cytochalasin D restores the NF-κB reporter activity. h, i) LPS increases the secretion of IL-6 and TNFα in RAW264.7 macrophages, which is inhibited by Resolvin-EVs and inhibition of Resolvin EV uptake restores the IL6 and TNFα secretion. Efferocytosis activity of THP-1-derived macrophages (Cyto Tell+ve) was measured by co-incubating with apoptotic neutrophils (PKH26+ve) and analyzed on BD Fortessa flow cytometer. j) PKH26+ve and Cyto Tell+ve population k) Co-culture with Resolvin-EVs enhances efferocytosis activity in macrophages (Cyto Tell+ve and PKH26+ve) compared to control EVs. a-i ****/####/x p ≤ 0.0001 for all comparisons.

### *In vivo* uptake of EVs by pulmonary cells

To determine *in vivo* EV uptake efficiency of different pulmonary cells, PKH26 (PE+) labeled EVs were delivered into Saline and LPS treated mice (4.0 mg/kg body weight i.n), after 1 h total pulmonary cells were isolated by collagenase digestion, stained with cell specific markers and analyzed by AMNIS Imagestream flow cytometer. Pulmonary macrophages, AEC, PMVEC and neutrophils showed significant differences in uptake efficiency (Figure 5a gating strategy, 5b relative PE+ve cell population). EV uptake in saline treated mouse lungs was maximum in macrophages (CD45+PE+F/480+ 20-23%), followed by PMVEC (CD45-PE+CD325-CD31+ 8-12%) and AEC (CD45-PE+CD31-CD326+ 4.6-5%). Interestingly, the EV uptake doubled in all the pulmonary cells from LPS-treated mice, close to 40% in macrophages; 35-42% in Neutrophils CD45+PE+F/480-Gr1+; 19-21% PMVEC and 16-20% AEC.

**Figure. 5.**
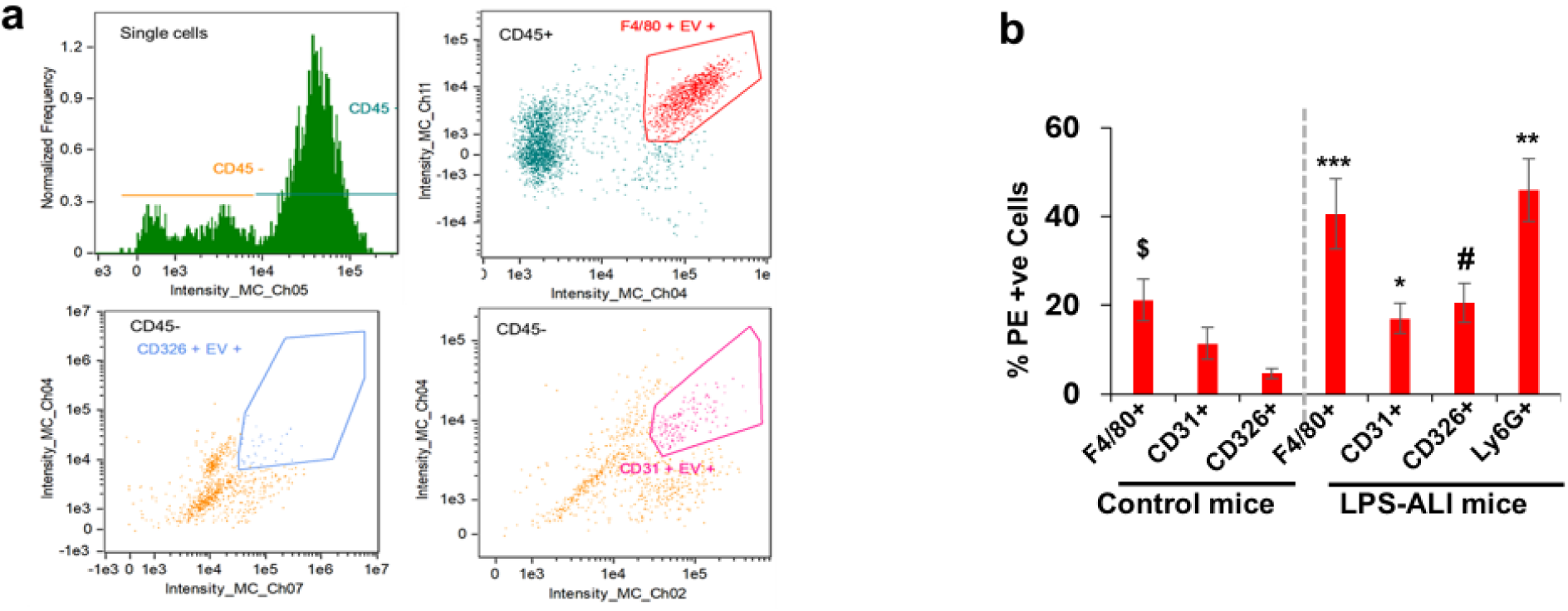
*Invivo* uptake of PKH26 labeled EVs by pulmonary cells. PKH26-labeled EVs were delivered (4 mg/kg, i.n.) into mice, and lung cells were isolated by collagenase digestion, stained with cell-specific markers, and analyzed using the AMNIS Imagestream flow cytometer. a) Representative gating-macrophages (CD45+F4/80+Ly6G-); neutrophils (CD45+F4/80-Ly6G+); epithelial cells (CD45-CD31-CD326+); endothelial cells (CD45-CD31+CD326-). b) Relative uptake of EVs by pulmonary cells from control and LPS-treated mice. */# p≤0.05; **/$$ p≤0.01; *** p≤0.001.

### Resolvin-EVs mediate active resolution of pulmonary inflammation in mouse models

We determined the *in vivo* efficacy of Resolvin-EVs in mitigating pulmonary inflammation in mice. C57BL6 mice were treated with a single dose of intranasal (i.n) LPS (4 mg/kg in 30 µL saline) and after 8h, either Control or Resolvin-EVs (4 mg/kg body weight, in 30 µL PBS) were delivered via i.n route. All the mice were euthanized after 16 h of EV treatment, BALF was collected and analyzed (Figure 6a-time line). LPS challenge increased influx of immune cells and neutrophils into alveolar space in Control EV treated mice (Figure 6b). Similarly, LPS significantly increased release of inflammatory cytokines (TNFα, IL6) and protein leakage into alveolar space (Figure 6c, 6d, 6e). In contrast, compared to control EVs, delivery of Resolvin-EVs into mice, 8 h after the LPS challenge dramatically decreased immune cell infiltration, TNFα, il-6, protein leak into BALF (Figure 6b-6e, Figure 7).

**Figure 6.**
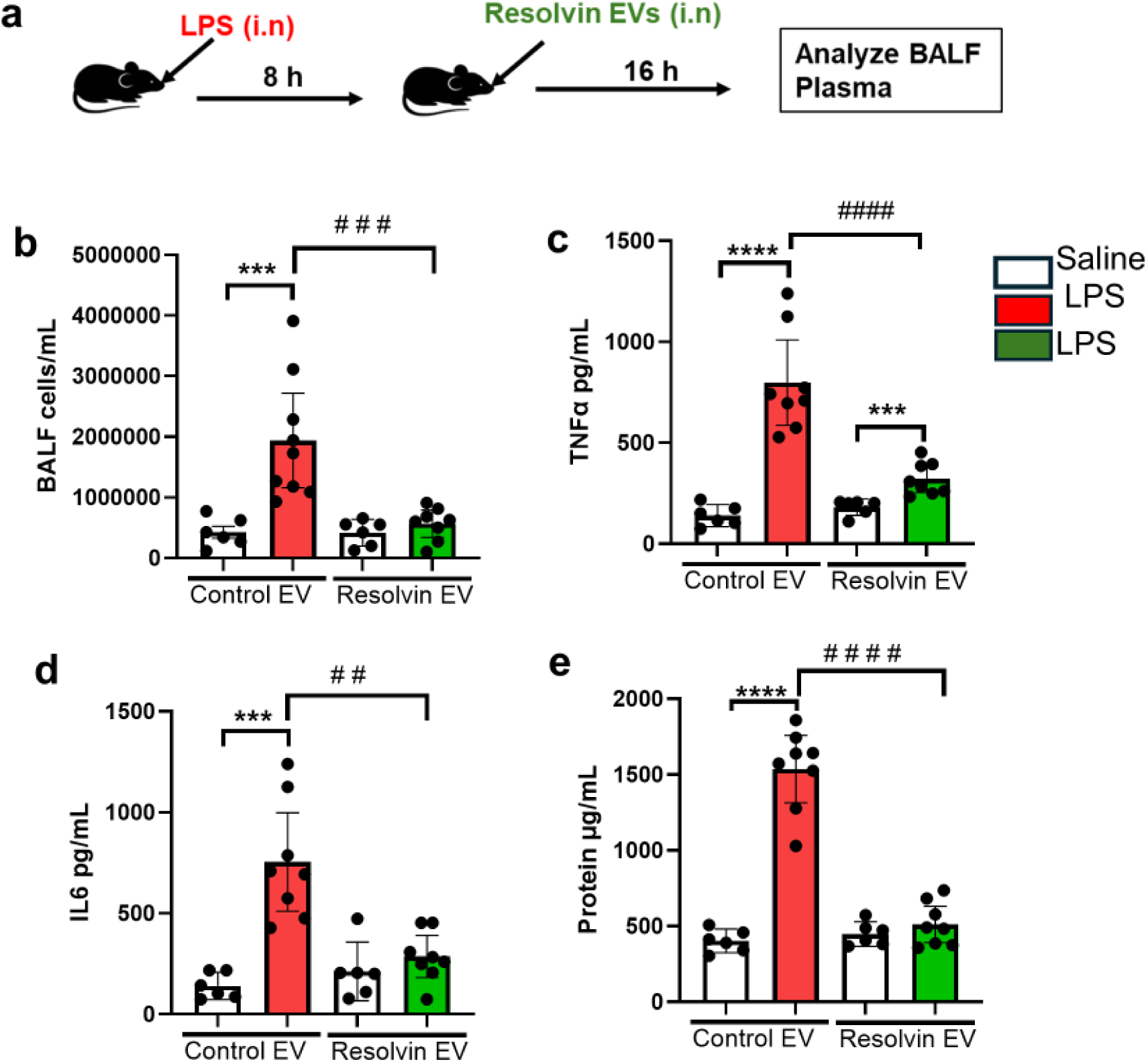
Resolvin-loaded EVs mitigate pulmonary inflammation in LPS-treated mice. **a)** C57BL6 mice were subjected to LPS-induced acute lung injury (4 mg/kg i.n). Eight hours after LPS treatment, Control EVs or Resolvin-EVs (4 mg/kg i.n) were administered. Sixteen hours of post-EV delivery, the mice were euthanized and analyzed. **b)** Total cells in BALF were separated by centrifugation and counted. Cell-depleted BALF was analyzed for: **c)** TNFα **d)** IL6 **e)** protein in alveolar fluid. **/## p≤0.01; ***/### p≤0.001; ****/#### p≤0.0001 n=6 control, n=8 in LPS groups.

**Figure 7.**
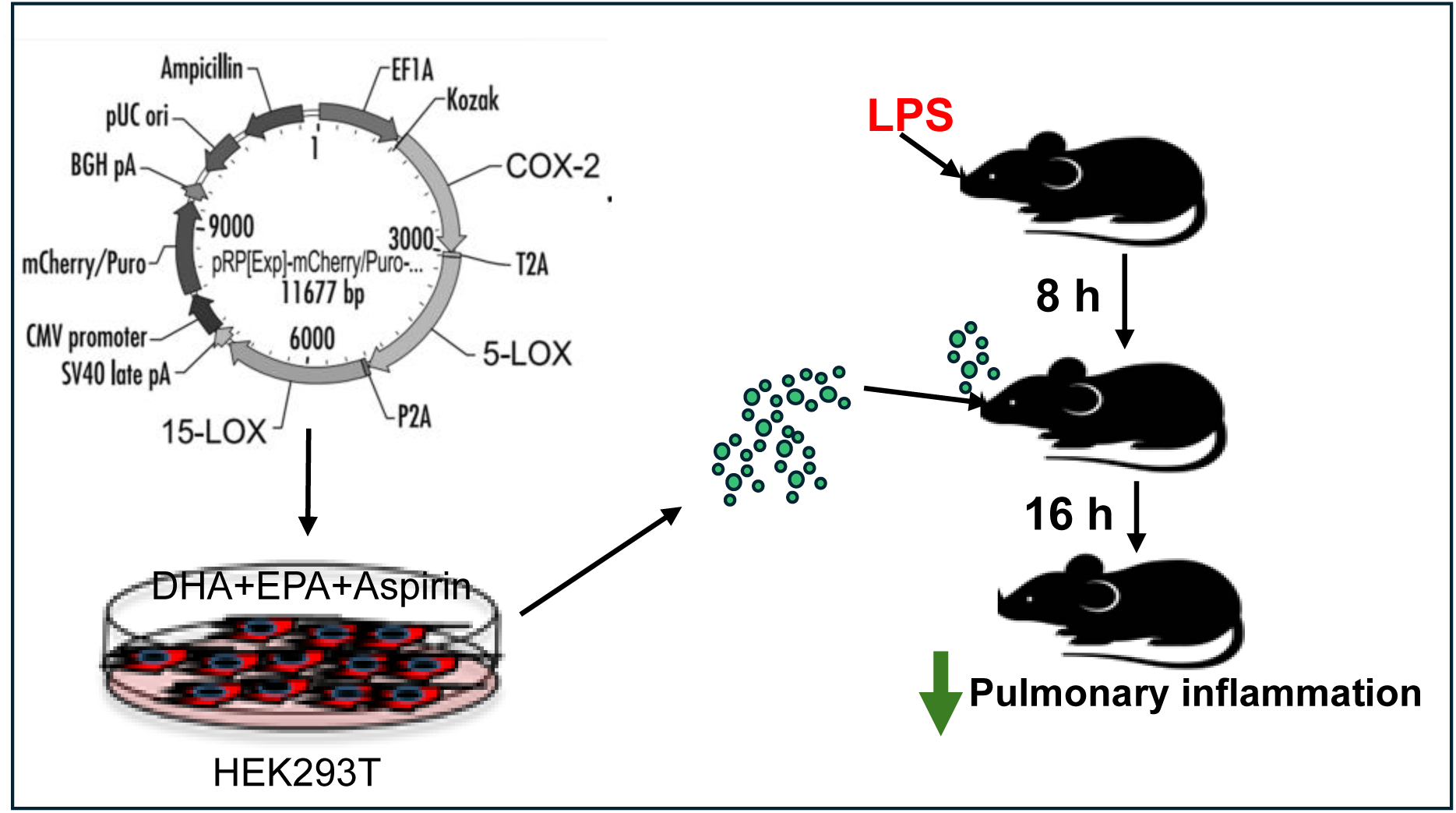
Schematic illustrating the potential of Resolvin EV targeted therapy for ALI. Resolvins are generated by mimicking transcellular and aspirin-triggered resolvin biosynthesis. The Resolvin loaded EVs actively mitigate pulmonary inflammation in LPS-treated mice by attenuating immune cell infiltration, cytokines release and protein leakage into alveolar space.

### *In vivo* toxicity of Resolvin-EVs

Mice were treated with saline or Resolvin-EVs (4 mg/kg i.n) and analyzed after 3 days. Plasma IL6 and TNFα levels in Resolvin-EV treated mice were comparable to saline-treated control mice (Table 2). Furthermore, IL6, TNFα and Lactate dehydrogenase (LDH) levels in BALF from Resolvin-EV treated mice were not elevated compared to saline-treated control mice. These findings emphasize the safety profile of Resolvin-EVs.

**Table 2.** Safety profile of Resolvin-EVs delivered via intranasal route into mice. Saline or Resolvin-EVs (4 mg/kg i.n) were administered into mice. 72 h post-EV delivery, the mice were euthanized and BALF analyzed for IL6, TNFα, immune cell infiltration and LDH. Similarly, plasma was analyzed for IL6 and TNFα. Measurements are mean ± SD, n=6 control, n=8 in Resolvin-EV groups.

|  | Analyte | Saline | Resolvin EV |
| --- | --- | --- | --- |
| BALF | IL6 pg/mL | 98±8.54 | 94±10.26 |
|  | TNFα pg/mL | 113±18.0 | 102±15.8 |
|  | BALF Cells | [8.9±0.98]x10<sup>5</sup> | [9.4±1.42]x10<sup>5</sup> |
|  | LDH | 4670± 616 RLU | 4309± 716 RLU |
| Plasma | IL6 pg/mL | 118± 11.22 | 98± 14.10 |
|  | TNFα pg/mL | 134.6± 21.57 | 121±18.28 |

## Discussion

Clinical studies have revealed a crucial role for various lipids in both the initiation and resolution of inflammation, particularly in critically ill patients and those suffering from acute respiratory distress syndrome (ARDS)^42–46^. In inflammatory contexts, prostaglandin E2 (PGE2) and leukotriene B4 (LTB4) function as neutrophilic and leukocyte chemoattractants. Notably, plasma analysis from ARDS non-survivors showed significantly elevated levels of the arachidonic acid metabolite LTB4^42^. Interestingly, peripheral blood from sepsis patients showed elevated levels of pro-inflammatory PGF2α, LTB4 in addition to SPMs, such as resolvin E1, resolvin D5, and 17R-protectin D1^43^. On the other hand, a study by Matthay et al. demonstrated that lung fluid from ARDS patients is enriched with LTD4, which correlates with pulmonary edema, while the levels of LTB4 and LTC4 remained unchanged^44^. Notably, critically ill and ARDS patients with uncomplicated recovery showed higher resolvin levels and lower leukotriene-to-resolvin ratios^45^. Collectively, these studies strongly support the role of lipid mediators as diagnostic and prognostic markers in ARDS and critical illness^43–46^.

Since their discovery, different members of SPMs have attracted considerable interest from researchers for their pivotal role in actively resolving inflammation^18, 21, 22^. Active resolution of inflammation represents a paradigm shift in understanding immune responses, focusing not on suppressing inflammation but on facilitating its active resolution, a process critical for maintaining tissue homeostasis and preventing both acute and chronic inflammatory diseases. Published data indicates that RvE1, RvE2 and RvE3 are biosynthesized from EPA by human PMNs via acetylated COX2 and 5-LOX, utilizing 18-HEPE as a precursor^18, 21, 47^.The biosynthesis of RvD1–6 occurs through two distinct pathways involving COX2, 5-LOX and 15-LOX enzymes, generating 17R- and 17S-series resolvins. The production of 17R-resolvins, or aspirin-triggered RvD (AT-RvD), is mediated by the conversion of DHA to a 17R-hydroperoxy intermediate by acetylated COX-2, whereas the formation of 17S-series resolvins (RvD1–6) is catalyzed by 15-LOX.

SPMs actively promote resolution through multiple mechanisms, including inhibition of neutrophil trans-endothelial and trans-epithelial migration, enhancement of macrophage efferocytosis to clear apoptotic cells, reduction of pro-inflammatory cytokine production/NF-kB activation, upregulation of the anti-inflammatory cytokine IL-10, and eventually establishing tissue homeostasis^11–15^. In addition to the pro-resolution effects of SPMs on immune cells, their role in mitigating the inflammatory phenotype of vascular and alveolar epithelial cells is well documented^48, 49^. SPMs not only reverse key cellular events in inflammatory conditions but also reduce the required dosage of antibiotics in infectious diseases^50, 51^. Conversely, reduced SPM production can exacerbate infection; macrophages from individuals with localized aggressive periodontitis contained lower levels of MaR1 compared to healthy individuals^52^. The cellular effects of SPM are mediated through interactions with specific G-protein-coupled receptors, such as ALX/FPR2 and ChemR23, expressed on immune cells, endothelial cells, and epithelial cells^53, 54^. Despite their potent bioactivity, SPMs are present in very low concentrations and have very short *in vivo* half-life. It has been shown that dietary supplementation with ω-3 PUFAs modulates immune responses, alleviating inflammatory diseases^55–58^. Although chemical synthesis methods of Resolvins are reported, they are not currently used as dietary supplements^59–61^.

We developed a novel approach to load EVs with a wide array of SPMs enriched at several thousand-fold higher levels than physiological concentrations, that we detected in mouse BALF. We previously demonstrated that BALF EVs from LPS-treated mice were packaged with high amounts of pro-inflammatory eicosanoids and significantly low levels of Lipoxin A4, Resolvin D1, and D6 (<20 pg/mg EV protein), suggesting the ability of pulmonary cells to package SPMs into EVs^35^. Interestingly, the enzymes COX-2, 5-LOX, and 15-LOX individually release pro-inflammatory eicosanoids from arachidonic acid^3,4^. However, these enzymes act in a coordinated fashion to produce SPMs from DHA and EPA^18, 19, 22^. We leveraged this overlapping biosynthetic pathway by artificially overexpressing COX and LOX enzymes in HEK293T cells and growing them in the presence of ω-3 fatty acids and aspirin to generate SPMs. Studies from our group and others demonstrate that lipids transported by EVs are stable due to their saturated lipid rich bilayer membrane. Furthermore, our study confirms that EVs enriched with resolvins accelerate the resolution of inflammation in both *in vitro* assays and *in vivo* mouse models. It is important to note that, unlike proteins and RNA, SPMs are structurally identical between mice and humans. Collectively, these features underscore the translational potential of Resolvin-EVs. Furthermore, studies indicate that EVs delivered via systemic route into mice are accumulated in liver and cleared rapidly, within few minutes^62^. We observed significant uptake of EVs by various pulmonary cells when administered through the i.n. route in mice. Notably, the Resolvin-EVs mitigated pulmonary inflammation in mice, when administered as post-injury treatment. The levels of TNFα, IL6 and LDH in BALF of Resolvin-EV treated mice were comparable to saline treated control mice, confirming the safety profile of Resolvin-EVs for pulmonary delivery in mice. In addition, HEK293 cells are commonly used for therapeutic protein production, show little to no expression of TLR4 receptor or MD-2 and CD14, suggesting a favorable safety profile for Resolvin-EVs generated from these cells. While most studies have evaluated a single pro-resolving lipid in *invitro* or rodent studies as an intervention, we have developed a novel strategy to load multiple SPMs into EVs. We demonstrate that the Resolvin-EVs to effectively mitigate pulmonary inflammation in mice as they contained multiple resolvin D and E members, that act by complimentary mechanisms.

Although we observed a higher number of Resolvin D members (D1, D2, D5 and D6), we detected only Resolvin E1 and E2 in Resolvin EVs. Furthermore, the concentration of Resolvin D1, D2, D5 and D6 is higher compared to Resolvin E1 and E2. This may represent a minor limitation of our approach, due to the differences in the substrate preferences of LOX biosynthetic enzymes. Nevertheless, given that Resolvin EVs are enriched with both resolvin D and E members exerting complementary cellular effects, we expect that Resolvin EVs will outperform the benefits of individual resolvins reported so far in the literature. Our studies, for the first time, introduced a novel scalable, cell-based Resolvin-loaded EV delivery platform that has potential to mitigate pulmonary inflammation and other acute inflammatory disorders.

## Methods

### Cell lines

Different cell lines used in the study were purchased from ATCC (HEK293T #CRL-3216; EA.hy926 #CRL-2922; HL60 #CCL-240; RAW 264.7 #TIB-71; THP-1 #TIB-202), human pulmonary microvascular endothelial cells (HPMVEC #H-6011 Cell biologics Inc, Chicago IL) and grown in appropriate ATCC/Cell Biologics Inc culture media, endothelial growth supplements, exosome depleted FBS and 1% penicillin/streptomycin. Serum free Opti-MEM was used for plasmid transfections.

### Fatty acid free BSA-complexed DHA and EPA preparation

DHA (Cayman Chemical # 90310) and EPA (Cayman Chemical # 90110) were complexed with fatty acid free BSA (Millipore Sigma #A8806) using a published protocol with minor modifications^63^. In brief, DHA and EPA (75 µM) separately dissolved in ethanol were gradually complexed with fatty acid-free BSA solution (150 µM) at 37^0^C for 2-4h and filtered through 50 kDa Millipore Sigma centrifugal filters. BSA-complexed fatty acid free DHA and EPA were added slowly to Opti-MEM, vortexed before adding to transfected HEK293T cells growing on cell culture dishes.

### Plasmid expression vectors

pcDNA3.1 Control, human pcDNA3.1_PTGS2, and pcDNA3.1_ALOX5 expression vectors were purchased from GenScript (Piscataway, NJ). pRP_Control and pRP_PTGS2_ALOX5_ALOX15 expression vectors were custom synthesized by VectorBuilder Inc. (Chicago, IL). PTGS2, ALOX5 and ALOX15 proteins were separated by P2A and T2A viral sequences for independent protein production.

### Generation of RvD1E1 EVs

6-8 x10^6^ HEK293T cells grown overnight in Opti-MEM were transfected with 10 µg each of human pcDNA3.1-COX2 and pcDNA3.1-5LOX plasmid vectors using nonlipid transfection reagent Fugene 6.0 (Promega, Madison, WI). After 24 hours of transfection, the medium was removed, cells washed and supplemented with fresh OptiMEM containing 10 µM each of DHA and EPA complexed with fatty acid free BSA, and 1 mM Asprin (Cayman, Detroit MI) and grown for another 24 h. After 24h, the culture medium was used for EV isolation and cells were lysed in 1X RIPA buffer for analyzing expression of human COX2 and 5-LOX proteins or used for total RNA isolation.

### Generation of Resolvin-EVs

6-8 x10^6^ HEK293T cells grown overnight in Opti-MEM were transfected with 20-25 µg each of pRP_Control or pRP_PTGS2_ALOX5_ALOX15 expression vectors. Resolvin-EVs were isolated from transfected cells grown in OptiMEM containing 10 µM each of DHA and EPA complexed with fatty acid free BSA, and 1 mM Asprin (Cayman, Detroit MI) and grown for another 24h-36 h. After 36 h, the culture medium was used for EV isolation and cells were lysed in 1X RIPA buffer for analyzing expression of human COX2 and 5-LOX proteins.

### Isolation of EVs from HEK293T cell supernatants

To isolate EVs, the cell culture medium was filtered through 0.22 µM Millipore syringe filters to remove microvesicles and the resulting filtrate was loaded on 20 mL Pierce 100 kDa cut off centrifugal filters (Thermofisher) and centrifuged at 4°C, 4000g for 60 min to remove BSA-complexed DHA, EPA and residual aspirin in culture medium. The concentrate is washed in 1X DPBS and resuspended in 100 μL of DPBS or saline for analysis. Based on the plasmid vector transfection and medium supplementation, 4 different groups of EVs were isolated: 1) pcDNA3.1 or pRP_Control vector transfected cells grown in the presence of solvent (0.001% ethanol) or DHA+EPA+1mM aspirin, labeled as vector Control EVs. 2) pcDNA3_COX2 and pcDNA_5LOX transfected cells, grown in the presence of 0.001% ethanol - referred as Control EVs 3) pcDNA3_COX2 and pcDNA_5LOX or custom synthesized, pRP_PTGS2_ALOX5_ALOX15 plasmid transfected cells, grown in presence of DHA+EPA+ASA mix-referred to as RvD1E1EVs or Resolvin-EVs which contain different resolvin members described in results section.

### Nanoparticle tracking analysis

The size of extracellular vesicles isolated from transfected HEK293T cells were determined as described previously, by Nano particle analysis using NanoSight at OSU Comprehensive Cancer Center Flow Cytometry facility^35, 36^. Mean size, particle numbers were compared between vesicles isolated from individual transfections, as described previously^35^.

### EV Proteomics

EVs were digested using sequencing grade Trypsin and analyzed at OSU Comprehensive Cancer Center-Proteomics Shared Resources (PSR) on the Thermo Orbitrap Fusion Mass Spectrometer. Data were searched against the database on MASCOT (version 2.7.0) using Proteome Discoverer (version 2.4.1.15) for the identification of the proteins. For protein IDs, false discovery rate was set at 1% for both peptide and protein level, a minimum of 2 unique peptides is required for valid protein identification. Mascot results were then compiled in scaffold (version 4.11.0) for quantitative and statistical analysis using label free spectral counting approach.

### RvD1E1 EVs lipidomics

LC-MS/MS analysis of RvD1E1 EVs was carried out at the National Jewish Health, Denver CO, lipidomics core facility as described previously^36^. Lipid extraction was performed with 0.1 N HCl to promote phase separation and enhance the recovery of free fatty acids and eicosanoids into the chloroform layer. LC-ESI-MS/MS was conducted with a Sciex 6500 QTRAP mass spectrometer, coupled to a Shimadzu Nexera-X2 UHPLC system. Free fatty acids and oxidized metabolites were separated on an Ascentis Express C18 column (2.1 × 50 mm, 2.7 μm). Gradient elution was carried out from solvent A (methanol:water:formic acid 30:70:0.1) to solvent B (methanol with 0.1% formic acid) at 0.65 mL/min. Deuterated arachidonic, eicosapentaenoic, and docosahexaenoic acids, along with prostaglandins, leukotrienes, and iso-prostanes (Cayman Chemicals, MI), were added during extraction (100 ng/sample for free fatty acids and 10 ng/sample for eicosanoids) for quantification using isotope dilution. Declustering potential and collision energy were optimized during infusion experiments.

### Resolvin-EV lipidomics

The control and Resolvin-EVs generated from pRP_Control and pRP_PTGS2_ALOX5_ALOX15 expression vector transfected cells were analyzed as described previously using liquid chromatorgraphy-mass spectrometry (LC-MS) at the lipidomics core facility, Wayne State University, Detroit^35^. In brief, SPM analysis was performed using High-Performance Liquid Chromatography (HPLC) and Liquid Chromatography-Mass Spectrometry (LC-MS) for fatty acyl quantification, as previously described^35^. Samples were extracted using C18 columns, redissolved in methanol, and analyzed by LC-MS. HPLC was conducted with a Prominence XR system (Shimadzu) and a Luna C18 column. The eluate was introduced into the ESI source of a QTRAP5500 mass spectrometer (SCIEX) in negative ion mode, monitored by Multiple Reaction Monitoring (MRM) for fatty acid detection. Mass spectra were recorded using Enhanced Product Ion (EPI) for peak identification, utilizing internal standards, MRM transitions, and retention time. Data were collected with Analyst 1.6.2 software, and MRM chromatograms quantified with MultiQuant software (from SCIEX). Internal standards were used to normalize different resolvins.

### Western blot analysis

Transfected HEK293T cells or EVs or human PMVEC cells were lysed in radio immune precipitation assay (RIPA) lysis buffer (Cell Signaling Technologies, Danvers, MA) with 1X protease inhibitor cocktail (ThermoFisher, Waltham, MA). Cell lysates containing 5-10 μg of total protein were electrophoresed on 4-20% SDS PAGE gradient gels, transferred on to PVDF membranes and immunoblotted with COX-2, 5-LOX, 15-LOX, CD9, Calnexin, VE-Cadherin antibodies and HRP-conjugated secondary antibodies, as described previously^35^. Expression levels of proteins were detected using a streptavidin-HRP chemiluminescence detection system (ThermoFisher, Waltham, MA).

### qRT-PCR analysis

Total RNA was isolated from pcDNA3.1_Control, pcDNA3.1_COX2, pcDNA3.1_5-LOX; pRP_Control or pRP_PTGS2_ALOX5_ALOX15 vector transfected cells and reverse transcribed using RevertAid RT reverse transcription kit (Thermofisher #K1691). Expression levels of COX2, 5-LOX and 15-LOX, and GAPDH in HEK293T cells and EVs were determined using gene specific primers (Supplementary material) as described previously^36^.

### Neutrophil-Endothelial cell adhesion assay

HL60 cells were differentiated to neutrophils and labeled with PKH26 cell linker as per the manufacturer’s instructions (Sigma Aldrich). EAhy926 cells were grown on 96 well-black polystyrene culture plates (Perkin Elmer) and treated with 50 ng/mL of TNFα for 8h. Following this, PKH26 labeled neutrophils were added to the EAhy926 cells and after 1h, 10 µg/mL of Control or RevD1E1EVs were added separately and incubated for additional 15 h. We have added equal amount of EV protein, regardless of EV number to avoid inconsistent delivery of cargo. After 16 h, the cells were washed three times with DPBS, and relative fluorescence (RFU) was measured on a plate reader at 530/570 nm. The cellular function of Control EVs, RvD1E1EVs or Resolvin-EVs was further confirmed by pretreating EAhy926 cells with EV uptake inhibitors, 50 µg/mL Nystatin and 10 µg/mL cytochalasin D separately in these assays. LDH activity in BALF was measured using LDH-Glo™ Cytotoxicity Assay kit (Promega #J2380) as per manufacturer’s instructions.

### PMVEC monolayer Barrier function measurements

PMVEC cells were grown to 100% confluence on collagen coated transwells and treated with 0.02% Thrombin for 30 min followed by control or RvD1E1 or Resolvin-EVs for 16h. FITC-Dextran was added to transwells at the time of EV addition and its flux to bottom wells was measured after overnight/16h incubation, as described previously^35, 64^.

### NF-kB luciferase reporter assay

Immortalized macrophages that express *Photinus* luciferase cDNA under the control of the nuclear factor NF-κB–dependent HIV-1 long terminal repeat promoter developed by our laboratory group were used to measure LPS-induced NF-kB activation^65, 66^. LPS-stimulated NF-kB reporter macrophages were co-incubated with Control or Resolvin-EVs for 3 hours. Cells were washed and lysed in cell lysis buffer and luciferase activity measured using luciferin as substrate (Promega # E152A).

### Mouse models of LPS-induced ALI

Wild type C57BL/6 (WT-Stock no. 000664) mice were purchased from Jackson Research Laboratories and were maintained at the pathogen free vivarium of The Ohio State University, Columbus, OH. Mice of 8-12-week age were used in the study. All the animal experiments were humanely conducted in accordance with protocols approved by the Ohio State University Institutional Animal Care and Use Committee (IACUC, #2013A00000105-R3). *Escherichia coli* lipopolysaccharide Serotype O55:B5 S-form, dissolved in sterile saline was delivered through intranasal insufflation in to anesthetized WT mice (4 mg/kg in 30 μL saline). Control mice received equal volume of saline. After 0 and 8 h after LPS treatment, mice were treated with Control or Resolvin-EVs (4 mg/kg in 30 μL saline). 16 h after the Resolvin-EV delivery, both the control and experimental mice were euthanized with ketamine/xylazine, trachea was cut open and BALF was collected by instilling 1 mL sterile saline three times and aspirating. Aspirated BALF was centrifuged at 300 x g to separate cells, and the supernatant was analyzed for TNFα, IL6, extravasated protein, as described previously^66^. Cytokine release was analyzed using R&D Systems ELISA kits for mouse TNF-α (catalog no. MTA00B) and IL-6 (catalog no. M6000B) following the protocols supplied by the manufacturer.

### Imagestream flow cytometry

To determine the in vivo uptake, EVs were labeled with PKH26 and purified on sucrose density gradient as per manufacturer’s instructions (#PKH26PCL-1KT, Sigma-Aldrich, St. Louis, MO) and delivered via i.n route into mice (4 mg/kg in 30 μL saline). 1 h after the EV delivery, both the control and experimental mice were euthanized with ketamine/xylazine and lungs harvested. Single cell suspension from total lungs were isolated by Collagenase/DNase, blocked with anti-mouse CD16/CD32 and stained with CD45, CD31, CD326, Ly6G and F4/80, as described previously^67^. Stained cells were washed and analyzed on the Amnis ImageStreamX Mark II Imaging Cytometer, at 60x magnification for all samples. A minimum of 10,000 cells per sample were analyzed. Data analysis was conducted with IDEAS software (Amnis Corporation).

### Statistical analysis

All data are expressed as mean ± SEM and were analyzed using Graphpad Prism 10.0. Differences between two experimental groups were compared with Student’s t test and comparisons among three or more groups were performed using ANOVA with a post hoc Bonferroni correction test, n=8 mice in each group processed separately for pulmonary inflammation measurements. p value <0.05 was considered significant. Heat maps and volcano plot were generated using Graphpad Prism 10.0.

## Supplementary Material

The DNA sequence of custom synthesized plasmid and PCR-Primes used in the study are included.

## Acknowledgements

The authors greatly acknowledge support from National Institutes of Health funding HL137224-JWC and MK; Ohio State University Clinical Translational Science Institute (OSU CTSI) #UM1TR004548 – MK; OSU flow cytometry facility is supported by P30CA016058. OSU Mass spectrometry and proteomics, Fusion Orbitrap instrument is supported by NIH Grant S10 OD018056. Wayne State University lipidomics core facility is supported by National Center for Research Resources (in part), NIH Grants S10RR027926 and S10OD032292 to Krishna Rao Maddipati (KRM).

## Author Contributions

Conceptualization, study design and data interpretation-MK; Manuscript writing and editing MK and JWC. Lipidomics analyses-EB and KRM; EV isolations, in vitro cell assays, and *in vivo* mouse experiments-JY and MK; Flow cytometry analyses-SC; interpretation of the data-MK, JWC, SRP and NLP. All the authors have approved the final submission.

## Data Availability Statement

All data generated or analyzed during this study are included in this article and its online supplementary material files. Further inquiries can be directed to the corresponding author.

