## Supplementary material for "Specialized Pro-Resolving Mediator loaded Extracellular Vesicles Mitigate Pulmonary Inflammation"

**Vector and sequence information:**

**Vector Name:**	pRP[Exp]-mCherry/Puro-EF1A >hPTGS2[NM_000963.4](co)(ns)*:T2A
:hALOX5[NM_000698.5](co)(ns)*:P2A :hALOX15[NM_001140.5](co)*  **[Abbreviated as pRP_PTGS2_ALOX5_ALOX15]**

**Vector Type:** Mammalian Gene Expression Vector

**Vector Size:** 11677 bp

**Promoter:** EF1A

**ORF:** hPTGS2[NM_000963.4](co)(ns)*, hALOX5[NM_000698.5](co)(ns)*, hALOX15[NM_001140.5](co)*

**Linker:** T2A, P2A

**Marker:** mCherry/Puro

**Plasmid Copy Number:** High

**Antibiotic Resistance:** Ampicillin

| **Name** | **Position** | **Size (bp)** | **Type** | **Description** | **Application notes** |
| --- | --- | --- | --- | --- | --- |
| EF1A | 22-1200 | 1179 | Promoter | Human eukaryotic translation elongation factor 1 α1 promoter | Strong promoter. |
| Kozak | 1225-1230 | 6 | Miscellaneo us | Kozak translation initiation sequence | Facilitates translation initiation of ATG start codon downstream of the Kozak sequence. |
| hPTGS2[NM_0  00963.4](co)  (ns)* | 1231-3042 | 1812 | CDS | None | None |
| T2A | 3043-3105 | 63 | Linker | Self-cleaving 2A peptide from Thosea asigna virus | Causes co-translational cleavage of the encoded polypeptide. Multiple proteins can be made from a polycistronic transcript containing multiple ORFs separated by 2A. P2A and T2A have higher cleavage efficiency compared to other 2As. |
| hALOX5[NM_0  00698.5](co)  (ns)* | 3106-5127 | 2022 | CDS | None | None |
| P2A | 5128-5193 | 66 | Linker | Self-cleaving 2A peptide  from Porcine teschovirus-1 | Causes co-translational cleavage of the encoded polypeptide. Multiple proteins can be made from a polycistronic transcript containing multiple ORFs separated by 2A. P2A and T2A have higher cleavage efficiency compared to other 2As. |
| hALOX15[NM_  001140.5](co)* | 5194-7182 | 1989 | CDS | None | None |
| Name | Position | Size (bp) | Type | Description | Application notes |
| SV40 late pA | 7227-7448 | 222 | PolyA_signal | Simian virus 40 late polyadenylation signal | Allows transcription termination and polyadenylation of mRNA transcribed by Pol II RNA polymerase. |
| CMV promoter | 7452-8039 | 588 | Promoter | Human cytomegalovirus immediate early enhancer/promoter | Strong promoter; may have variable strength in some cell types. |
| mCherry/Puro | 8071-9378 | 1308 | CDS | mCherry fused with Puro | Allows cells to be visualized by red fluorescence and resistant to puromycin. |
| BGH pA | 9422-9646 | 225 | PolyA_signal | Bovine growth hormone polyadenylation signal | Allows transcription termination and polyadenylation of mRNA transcribed by Pol II RNA polymerase. |
| pUC ori | complement (9842-  10430) | 589 | Rep_origin | pUC origin of replication | Facilitates plasmid replication in E. coli; regulates high-copy plasmid number (500700). |
| AmpiciIIin | complement (10601-  11461) | 861 | CDS | AmpiciIIin resistance gene | Allows E. coli to be resistant to ampiciIIin. |

**hPTGS2 [NM_000963.4] (COX2) codon optimized nucleotide sequence**

ATGCTGGCCA GGGCCCTGCT CCTGTGCGCC GTGTTGGCCC TGTCACACAC CGCCAATCCC TGTTGCTCAC ACCCCTGCCA GAACAGAGGC GTCTGTATGA GCGTGGGCTT CGACCAGTAC AAGTGCGATT GCACTAGAAC GGGCTTTTAC GGAGAGAACT GTAGCACTCC CGAGTTCCTG ACTCGCATCA AACTGTTCCT GAAGCCTACA CCCAATACCG TGCACTATAT TCTGACACAC TTTAAGGGCT TCTGGAATGT GGTGAACAAT ATCCCATTTC TGAGAAATGC TATCATGAGC TACGTGCTGA CTAGTAGGTC TCACTTGATC GACTCCCCAC CTACTTATAA TGCCGACTAT GGCTACAAGT CCTGGGAAGC CTTCTCCAAT CTGAGTTACT ATACCAGGGC TCTGCCCCCT GTGCCAGACG ATTGCCCTAC ACCTCTGGGC GTCAAGGGGA AGAAGCAGCT GCCCGACTCC AATGAGATTG TGGAGAAACT GCTCCTCAGG CGGAAGTTTA TTCCTGACCC TCAGGGAAGC AACATGATGT TCGCCTTCTT CGCCCAGCAC TTCACCCACC AGTTCTTTAA AACAGACCAT AAGCGCGGCC CTGCCTTCAC CAACGGCCTG GGCCACGGCG TCGACCTAAA TCATATCTAC GGCGAAACCC TGGCCCGGCA GAGGAAGCTG AGGCTGTTCA AAGACGGCAA GATGAAGTAC CAGATTATTG ACGGCGAGAT GTACCCTCCT ACCGTGAAGG ACACCCAGGC AGAGATGATC TATCCCCCCC AGGTGCCTGA GCACCTGAGG TTCGCTGTGG GGCAGGAAGT GTTCGGCCTG GTGCCCGGGC TGATGATGTA CGCAACCATC TGGCTGCGGG AACACAATAG AGTGTGCGAT GTCCTGAAAC AGGAGCACCC CGAGTGGGGC GACGAGCAGC TGTTCCAGAC ATCTCGGCTG ATCCTCATTG GTGAGACTAT CAAGATCGTG ATCGAGGACT ACGTGCAGCA CCTGAGCGGA TACCACTTTA AGCTGAAATT TGATCCTGAA CTGCTGTTCA ACAAGCAGTT TCAGTACCAG AACAGGATTG CCGCCGAGTT TAACACCCTG TACCACTGGC ACCCCCTGCT GCCAGACACC TTCCAGATCC ACGACCAGAA ATATAATTAT CAGCAGTTCA TCTACAACAA CTCAATCCTG CTGGAACACG GCATCACACA GTTCGTGGAG AGTTTCACCA GACAGATCGC CGGCAGGGTG GCCGGGGGCA GGAATGTGCC CCCCGCCGTG CAGAAAGTGT CCCAGGCCTC TATCGACCAG TCTAGGCAGA TGAAGTACCA GTCCTTCAAC GAGTACAGAA AGCGGTTCAT GCTGAAGCCC TACGAGAGCT TCGAGGAACT GACCGGAGAA AAGGAGATGA GCGCCGAGCT GGAGGCCCTG TATGGAGACA TTGACGCCGT GGAGCTGTAC CCCGCTCTGC TGGTGGAGAA ACCACGCCCC GACGCCATCT TTGGGGAGAC CATGGTGGAA GTGGGAGCCC CCTTTTCCCT GAAGGGCCTG ATGGGCAACG TGATCTGCAG CCCCGCCTAT TGGAAACCTT CCACCTTCGG CGGCGAGGTG GGATTCCAGA TCATCAACAC CGCCTCTATC CAGTCCCTGA TCTGCAATAA TGTGAAAGGC TGTCCCTTCA CCAGCTTCAG CGTGCCCGAT CCCGAGCTGA TCAAAACCGT GACCATCAAT GCCTCCTCCT CAAGAAGCGG ACTGGACGAC ATTAACCCCA CCGTGCTGCT GAAGGAGCGG AGTACCGAAC TG

**hPTGS2 [NM_000963.4] protein sequence**

MLARALLLCAVLALSHTANPCCSHPCQNRGVCMSVGFDQYKCDCTRTGFYGENCSTPEFLTRIKLFLKPTPNTVHYILTHFKGFWNVVNNIPFLRNAIMSYVLTSRSHLIDSPPTYNADYGYKSWEAFSNLSYYTRALPPVPDDCPTPLGVKGKKQLPDSNEIVEKLLLRRKFIPDPQGSNMMFAFFAQHFTHQFFKTDHKRGPAFTNGLGHGVDLNHIYGETLARQRKLRLFKDGKMKYQIIDGEMYPPTVKDTQAEMIYPPQVPEHLRFAVGQEVFGLVPGLMMYATIWLREHNRVCDVLKQEHPEWGDEQLFQTSRLILIGETIKIVIEDYVQHLSGYHFKLKFDPELLFNKQFQYQNRIAAEFNTLYHWHPLLPDTFQIHDQKYNYQQFIYNNSILLEHGITQFVESFTRQIAGRVAGGRNVPPAVQKVSQASIDQSRQMKYQSFNEYRKRFMLKPYESFEELTGEKEMSAELEALYGDIDAVELYPALLVEKPRPDAIFGETMVEVGAPFSLKGLMGNVICSPAYWKPSTFGGEVGFQIINTASIQSLICNNVKGCPFTSFSVPDPELIKTVTINASSSRSGLDDINPTVLLKERSTEL

**hALOX5 [NM_000698.5] codon optimized nucleotide sequence**

ATGCC CTCCTACACC GTCACCGTGG CCACTGGCTC CCAGTGGTTC GCCGGCACCG ATGACTACAT CTACCTGTCC CTGGTGGGCA GCGCCGGCTG CAGCGAGAAG CACCTGCTGG ATAAGCCCTT CTACAACGAT TTCGAGAGGG GCGCCGTGGA CAGTTATGAT GTGACTGTGG ACGAGGAACT GGGCGAGATC CAGCTGGTCC GCATCGAGAA AAGGAAATAC TGGCTCAACG ACGATTGGTA CCTGAAGTAT ATCACCCTGA AAACACCACA CGGAGACTAT ATTGAGTTCC CATGCTACAG GTGGATCACC GGCGATGTGG AGGTCGTGCT GCGGGATGGA AGGGCCAAGC TGGCCAGGGA TGATCAGATC CACATCCTGA AGCAGCACCG CAGGAAAGAG CTGGAGACCC GGCAGAAGCA ATATAGGTGG ATGGAGTGGA ATCCCGGCTT CCCTCTGTCC ATCGACGCCA AGTGCCACAA AGATCTGCCA AGGGACATAC AGTTTGACAG CGAGAAAGGC GTGGACTTCG TGCTCAACTA CAGCAAGGCA ATGGAGAACC TCTTTATCAA TAGATTTATG CACATGTTTC AAAGCTCCTG GAACGATTTT GCCGACTTCG AAAAAATCTT CGTGAAGATC AGCAATACCA TCTCCGAAAG AGTGATGAAT CACTGGCAGG AGGACCTGAT GTTCGGCTAC CAGTTCCTGA ACGGCTGCAA TCCAGTTCTG ATCAGACGGT GTACTGAGCT GCCTGAGAAG CTGCCCGTGA CAACAGAGAT GGTCGAGTGC TCCCTGGAAC GGCAGCTGTC CCTGGAGCAG GAGGTGCAGC AGGGCAACAT CTTCATCGTG GACTTCGAGC TGCTGGACGG AATTGACGCC AACAAAACCG ACCCCTGTAC CCTGCAGTTC CTGGCTGCCC CAATTTGCCT CCTGTACAAA AATCTGGCCA ATAAGATCGT GCCTATCGCC ATTCAGCTGA ACCAGATTCC CGGAGACGAG AATCCCATCT TTCTGCCTTC TGACGCCAAG TACGACTGGC TGCTGGCTAA GATCTGGGTG AGATCTAGCG ACTTTCACGT GCATCAGACC ATCACCCACC TCCTGAGAAC CCACCTGGTC AGCGAGGTGT TCGGGATTGC CATGTACCGG CAGCTACCCG CCGTGCACCC CATCTTCAAG CTGCTGGTGG CCCACGTGAG GTTTACAATT GCCATCAATA CGAAGGCACG AGAACAGCTG ATCTGCGAGT GCGGCCTGTT CGACAAGGCA AACGCCACCG GCGGCGGCGG ACACGTGCAG ATGGTCCAGC GAGCCATGAA AGACCTGACA TATGCTAGCC TGTGTTTTCC CGAAGCCATC AAGGCCAGAG GCATGGAGTC AAAAGAGGAC ATCCCCTACT ATTTCTATCG GGATGATGGC CTGCTGGTGT GGGAGGCCAT CCGGACATTC ACAGCCGAGG TGGTGGACAT CTATTATGAG GGGGACCAGG TGGTGGAGGA GGATCCCGAA CTGCAGGACT TCGTGAACGA CGTGTACGTG TACGGCATGA GGGGAAGGAA GTCCTCCGGA TTCCCAAAAA GCGTGAAGAG CCGCGAGCAG CTGAGCGAGT ACCTGACAGT GGTGATTTTT ACTGCCAGCG CCCAGCACGC CGCTGTGAAC TTCGGCCAGT ACGACTGGTG CAGCTGGATT CCCAATGCCC CACCCACAAT GCGCGCCCCC CCCCCAACCG CCAAGGGCGT GGTGACAATT GAGCAGATCG TGGACACACT GCCTGACCGC GGCAGGAGCT GTTGGCATCT GGGCGCCGTG TGGGCCCTGA GCCAGTTTCA GGAGAACGAG CTGTTCCTGG GGATGTATCC TGAGGAGCAT TTCATCGAGA AGCCCGTGAA GGAGGCCATG GCCCGCTTCA GAAAAAATCT GGAGGCCATC GTGAGCGTGA TCGCCGAGCG CAATAAAAAG AAGCAGCTGC CCTACTATTA CCTGTCACCC GATAGGATCC CAAACTCTGT GGCCATT

**hALOX5[NM_000698.5 protein sequence**

MPSYTVTVATGSQWFAGTDDYIYLSLVGSAGCSEKHLLDKPFYNDFERGAVDSYDVTVDEELGEIQLVRIEKRKYWLNDDWYLKYITLKTPHGDYIEFPCYRWITGDVEVVLRDGRAKLARDDQIHILKQHRRKELETRQKQYRWMEWNPGFPLSIDAKCHKDLPRDIQFDSEKGVDFVLNYSKAMENLFINRFMHMFQSSWNDFADFEKIFVKISNTISERVMNHWQEDLMFGYQFLNGCNPVLIRRCTELPEKLPVTTEMVECSLERQLSLEQEVQQGNIFIVDFELLDGIDANKTDPCTLQFLAAPICLLYKNLANKIVPIAIQLNQIPGDENPIFLPSDAKYDWLLAKIWVRSSDFHVHQTITHLLRTHLVSEVFGIAMYRQLPAVHPIFKLLVAHVRFTIAINTKAREQLICECGLFDKANATGGGGHVQMVQRAMKDLTYASLCFPEAIKARGMESKEDIPYYFYRDDGLLVWEAIRTFTAEVVDIYYEGDQVVEEDPELQDFVNDVYVYGMRGRKSSGFPKSVKSREQLSEYLTVVIFTASAQHAAVNFGQYDWCSWIPNAPPTMRAPPPTAKGVVTIEQIVDTLPDRGRSCWHLGAVWALSQFQENELFLGMYPEEHFIEKPVKEAMARFRKNLEAIVSVIAERNKKKQLPYYYLSPDRIPNSVAI

**hALOX15[NM_001140.5 codon optimized nucleotide sequence**

ATGGGCT TGTACCGGAT CAGGGTGAGT ACCGGCGCCA GCCTGTATGC TGGGAGCAAT AATCAGGTGC AGCTGTGGCT GGTGGGCCAG CACGGAGAGG CCGCCCTGGG CAAGCGCCTG TGGCCAGCTA GAGGCAAGGA AACAGAACTG AAAGTGGAGG TGCCCGAGTA TCTGGGCCCC CTGCTGTTCG TGAAACTCAG AAAAAGACAC CTGCTGAAGG ACGACGCCTG GTTCTGTAAT TGGATCTCTG TGCAGGGCCC CGGCGCCGGT GACGAGGTGA GATTCCCCTG CTACAGGTGG GTGGAGGGGA ATGGCGTGCT TTCCCTGCCC GAGGGGACAG GACGGACTGT GGGAGAGGAC CCACAGGGGC TGTTTCAGAA GCACCGGGAG GAGGAACTGG AAGAGCGCAG AAAGCTGTAC AGATGGGGCA ACTGGAAGGA CGGCCTGATC CTGAACATGG CTGGGGCCAA ACTGTACGAT CTGCCAGTGG ACGAGCGCTT TCTGGAAGAC AAAAGAGTTG ATTTCGAAGT GAGTCTGGCC AAGGGCCTGG CCGACCTGGC CATTAAAGAT TCCCTGAATG TGCTGACCTG CTGGAAAGAC CTGGATGATT TCAATCGGAT TTTCTGGTGT GGGCAGAGTA AGCTGGCAGA GAGGGTGAGG GATAGCTGGA AGGAGGACGC ACTGTTTGGG TATCAGTTCC TGAATGGCGC CAACCCTGTA GTGCTGAGAC GGAGCGCCCA TCTGCCCGCT AGACTGGTGT TTCCCCCTGG CATGGAGGAA CTGCAGGCCC AGCTGGAGAA GGAGCTGGAG GGCGGAACCC TGTTCGAGGC CGATTTTAGC CTGCTGGACG GCATCAAGGC CAATGTGATT CTGTGCTCTC AGCAGCACCT CGCCGCCCCT CTGGTGATGC TGAAGCTGCA GCCCGATGGT AAGCTGCTGC CCATGGTGAT CCAGCTGCAG CTGCCTCGCA CCGGCAGCCC ACCTCCCCCA CTGTTCCTGC CTACCGATCC CCCTATGGCC TGGCTGCTGG CCAAGTGCTG GGTGCGGAGC AGCGATTTCC AGCTGCACGA GCTGCAGTCA CACCTGCTGA GAGGCCACCT GATGGCCGAA GTGATCGTGG TGGCAACCAT GAGGTGTCTC CCCAGCATTC ACCCCATTTT TAAGCTGATC ATCCCACACC TGCGGTACAC CCTGGAGATC AATGTGCGAG CCAGAACAGG CCTGGTCAGC GACATGGGAA TCTTCGACCA GATTATGTCC ACCGGGGGGG GCGGTCACGT GCAGCTGCTG AAACAGGCTG GGGCTTTCCT GACTTACAGT TCCTTCTGTC CACCTGACGA CCTGGCCGAT AGGGGGCTGC TGGGAGTCAA GAGCTCCTTC TACGCTCAGG ACGCCCTGCG GCTGTGGGAG ATCATCTACC GGTACGTGGA GGGCATCGTG AGCCTGCACT ACAAGACTGA CGTGGCCGTG AAGGACGATC CTGAGCTGCA GACCTGGTGC AGAGAGATCA CCGAGATCGG CCTGCAGGGG GCCCAGGATA GAGGCTTTCC CGTGTCACTG CAGGCCAGGG ATCAGGTGTG CCACTTTGTG ACCATGTGCA TTTTTACATG CACAGGCCAG CACGCATCCG TGCACCTGGG CCAGCTTGAT TGGTATAGCT GGGTGCCCAA CGCTCCTTGC ACCATGCGCC TGCCTCCCCC CACCACCAAA GATGCCACCT TGGAGACCGT GATGGCAACT CTGCCCAACT TTCACCAGGC CAGCCTGCAG ATGAGCATCA CTTGGCAGCT CGGGCGCAGA CAGCCCGTAA TGGTGGCTGT GGGACAGCAC GAGGAGGAGT ACTTCAGCGG CCCAGAACCT AAGGCCGTGC TGAAGAAGTT TCGCGAGGAG CTGGCCGCCC TGGACAAAGA GATCGAGATC AGGAATGCAA AGCTGGATAT GCCCTATGAA TATCTGAGGC CCAGCGTGGT GGAGAACAGC GTGGCCATCT GA

**hALOX15[NM_001140.5 protein sequence**

MGLYRIRVSTGASLYAGSNNQVQLWLVGQHGEAALGKRLWPARGKETELKVEVPEYLGPLLFVKLRKRHLLKDDAWFCNWISVQGPGAGDEVRFPCYRWVEGNGVLSLPEGTGRTVGEDPQGLFQKHREEELEERRKLYRWGNWKDGLILNMAGAKLYDLPVDERFLEDKRVDFEVSLAKGLADLAIKDSLNVLTCWKDLDDFNRIFWCGQSKLAERVRDSWKEDALFGYQFLNGANPVVLRRSAHLPARLVFPPGMEELQAQLEKELEGGTLFEADFSLLDGIKANVILCSQQHLAAPLVMLKLQPDGKLLPMVIQLQLPRTGSPPPPLFLPTDPPMAWLLAKCWVRSSDFQLHELQSHLLRGHLMAEVIVVATMRCLPSIHPIFKLIIPHLRYTLEINVRARTGLVSDMGIFDQIMSTGGGGHVQLLKQAGAFLTYSSFCPPDDLADRGLLGVKSSFYAQDALRLWEIIYRYVEGIVSLHYKTDVAVKDDPELQTWCREITEIGLQGAQDRGFPVSLQARDQVCHFVTMCIFTCTGQHASVHLGQLDWYSWVPNAPCTMRLPPPTTKDATLETVMATLPNFHQASLQMSITWQLGRRQPVMVAVGQHEEEYFSGPEPKAVLKKFREELAALDKEIEIRNAKLDMPYEYLRPSVVENSVAI

**qRT-PCR Primer sequences:**

**COX2 (PTGS2)**

Forward 5' - CGG TGA AAC TCT GGC TAG ACA G - 3'

Reverse 5' - GCA AAC CGT AGA TGC TCA GGG A - 3'

**5-LOX (ALOX5)**

Forward 5' - GGA GAA CCT GTT CAT CAA CCG C - 3'

Reverse 5' - CAG GTC TTC CTG CCA GTG ATT C - 3'

**15-LOX (ALOX15)**

Forward 5' - ACC TTC CTG CTC GCC TAG TGT T - 3'

Reverse 5' - GGC TAC AGA GAA TGA CGT TGG C - 3'

**GAPDH**

Forward 5' - GAA GGT GAA GGT CGG AGTC - 3'

Reverse 5' - GAA ATG GTG ATG GGA TTG - 3'

**RPLP0**

Forward 5' - GGA GAA ACT GCT GCC TCA TAT C- 3'

Forward 5' - CAG CAG CTG GCA CCT TAT T- 3'
